# Label-free imaging of intracellular structures in living mammalian cells via external apodization phase-contrast microscopy

**DOI:** 10.1101/2024.03.01.582671

**Authors:** Hiroshi Ohno, Takenori Nishimura, Kenta Kainoh, Yoshitaka Ohashi, Naoko Onodera, Mayuko Kano, Lay Nurhana Sari, Masato Masuda, Yoshiaki Tamura, Yusuke Hayashi, Yusuke Yamamoto, Shin-Ichiro Takahashi, Yuta Mishima, Yosuke Yoneyama, Yoshinori Takeuchi, Motohiro Sekiya, Takashi Matsuzaka, Takafumi Miyamoto, Hitoshi Shimano

## Abstract

Developing techniques to visualize intracellular structures, which influence the spatiotemporal functionality of biomolecules, is essential for elucidating mechanisms governing cellular behavior. In this study, we demonstrate that label-free external apodization phase-contrast (ExAPC) microscopy serves as a valuable tool for the simultaneous observation of various intracellular structures with high spatiotemporal resolution, while successfully mitigating halo artifacts. Additionally, through quantitative analysis of images obtained by combining ExAPC microscopy with fluorescence microscopy, we identified distinct heterogeneities in biomolecular condensates, lipid droplets, and mitochondria. Our findings highlight the potential of ExAPC microscopy to provide detailed insights into alterations in intracellular structures associated with diverse cellular processes, corroborating the existing knowledge and potentially contributing to the discovery of novel cellular mechanisms.

## Introduction

Cells harbor intracellular structures called organelles that perform many diverse functions. The characteristic dimensions, quantities, and spatial arrangements of each organelle lead to the overall morphological attributes of the cell, termed cellular organization^1^. Many researchers have suggested that cellular organization, in concert with the functional attributes of cellular organelles, plays a pivotal role in determining diverse cellular behaviors^2–5^.

To understand the significance of cellular organization and its influence on various cellular behaviors, it is essential to visualize and clarify the intricate features of intracellular structures. Fluorescence imaging has become a fundamental tool for this purpose^6,7^. This technique enables detailed visualization of intracellular structures by employing fluorescent markers that specifically localize to those structures. Advances in molecular genetics have made fluorescent labeling more accessible, offering advantages such as speed, specificity, and applicability across various model organisms. However, fluorescence imaging has limitations, including photobleaching and phototoxicity^8^. In response, recent attention has shifted toward label-free imaging techniques, which exploit the optical contrast generated by light interactions with cells^9^. This approach avoids the need for labeling and generates cellular images in a state closer to the native environment.

Several innovative label-free imaging techniques have been developed^10–12^, including apodized phase contrast microscopy^13,14^. In the apodized phase contrast microscopy, the sample is positioned in the front focal plane of the objective lens and illuminated through an annular aperture. The incident light passes through an apodized phase ring surrounded by two light-attenuating rings, inducing a phase shift. The diffracted light passing through these attenuating rings is minimized, reducing halo artifacts^15^. Although commercially available objective lenses with an apodized phase plate exist, high-magnification lenses suitable for observing intracellular structures are not accessible. Nevertheless, the value of advanced visualization techniques lies in ensuring their accessibility to the broader scientific community. To address this gap, we employed an external apodized phase contrast (ExAPC) microscopy method, in which an apodized phase plate is placed outside the objective lens (Figure 1A). This approach overcomes the inherent limitations of commercially available lenses and facilitates the simultaneous imaging of multiple organelles while significantly reducing halo artifacts compared to conventional phase-contrast microscopy (Figure 1B).

**Figure 1.**
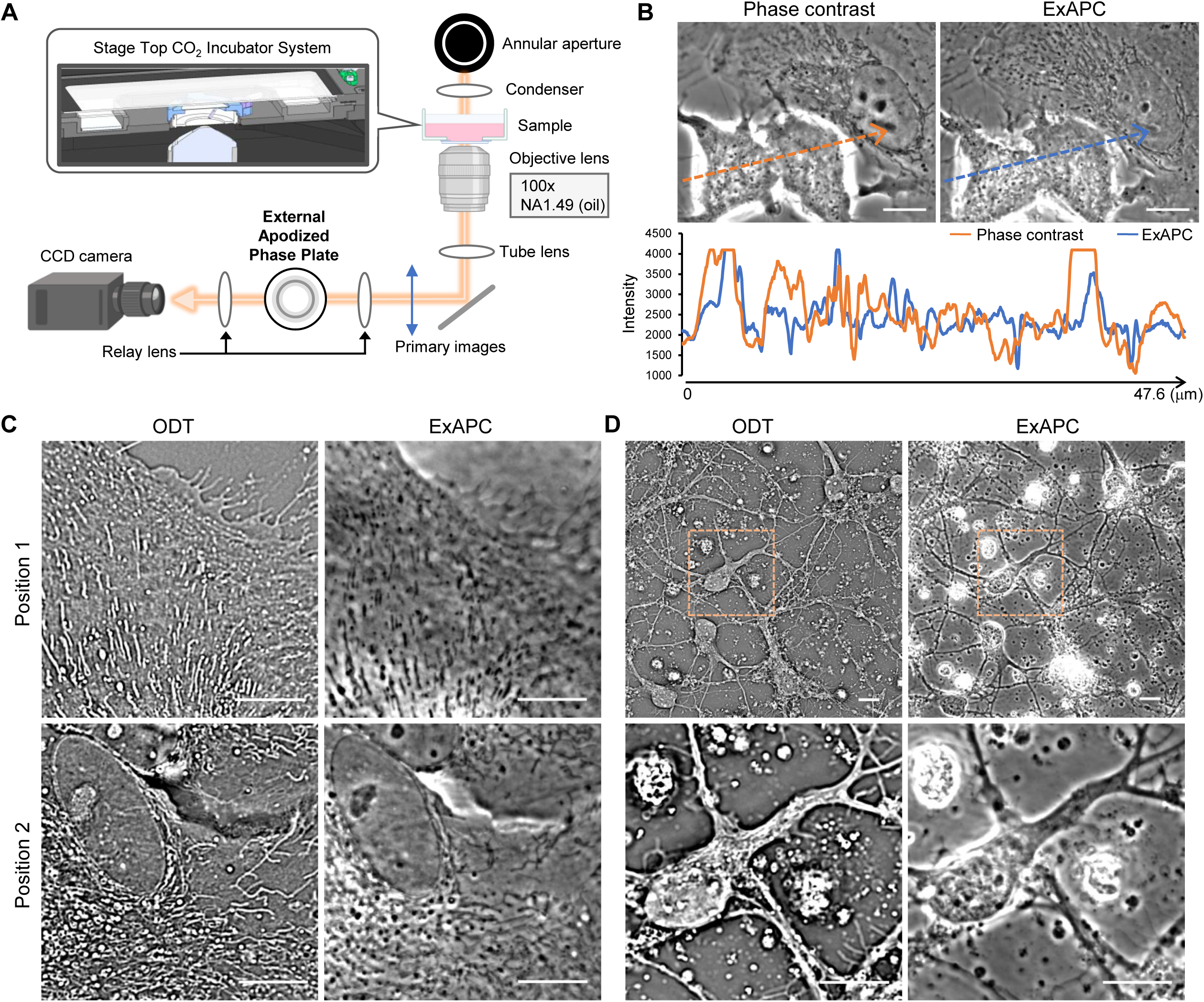
Visualization of cellular organization utilizing ExAPC microscopy. (A) Schematic representation of the ExAPC microscope used in this study. (B) Phase-contrast and ExAPC microscopy images of live A549 cells. (C) ODT and ExAPC microscopy images of live primary astrocytes. (D) ODT and ExAPC microscopy images of live primary neurons. All scale bar: 10 μm.

In this study, we demonstrated the utility of the ExAPC microscopy technique for observing cellular organization. Furthermore, through a detailed examination of biomolecular condensates, lipid droplets, and mitochondria that constitute cellular organization, we revealed the diversity in their behaviors and characteristics.

## Results

### Comparison of ExAPC microscopy and ODT for imaging cellular organization

ExAPC microscopy is a two-dimensional phase-imaging technique, whereas optical diffraction tomography (ODT) as a technique to visualize intracellular structures enables three-dimensional phase imaging and has recently garnered attention. A comparison between ExAPC microscopy and ODT showed that both techniques provide comparable results for visualizing the morphology of intracellular structures (Figure 1C). A major difference between these techniques lies in their temporal resolution: ExAPC microscopy achieves sub-second imaging intervals (Video 1), whereas obtaining comparable spatial and temporal resolutions with ODT, which requires Z-axis imaging, remains challenging^16,17^. Despite this, ODT offer the advantage of imaging thicker specimens, a task that presents limitations for ExAPC microscopy (Figure 1D). These results indicate that the choice between ExAPC microscopy and ODT should depend on experimental objectives, as both modalities offer complementary advantages in advancing our understanding of cellular organization.

### Visualization of cellular organization in live cells using ExAPC microscopy

The ExAPC microscopy system facilitates the label-free visualization of diverse intracellular structures within living cells. Nevertheless, the unambiguous identification of these structures based solely on the images obtained remains a substantial challenge. To overcome this limitation, we combined ExAPC microscopy with fluorescence imaging to accurately identify organelles detectable using the ExAPC system. Structures such as nuclei, nucleosomes, and mitochondria, characterized by refractive indices distinct from the cytoplasm^17^, were effectively visualized through ExAPC microscopy (Figure 2A, B). Additionally, actin filaments were discernible using this approach (Figure 2C). While the endoplasmic reticulum (ER) was difficult to detect in proximity to the nucleus, peripheral regions of the cell revealed structures partially colocalizing with ER-specific markers (Figure 2D). Furthermore, numerous vesicular structures, positively stained with markers for endosomes and lysosomes, were identified using ExAPC microscopy system (Figure. 2E, F). The average size of these vesicles was calculated to be 0.177 ± 0.168 μm² (n = 1,239), consistent with previously reported dimensions for these organelles^18,19^. Therefore, these vesicular structures were designated as “endosome- and lysosome-like structures (ELL)”. Conversely, the Golgi apparatus, which exhibits minimal refractive index contrast with the cytoplasm, remained challenging to detect using ExAPC microscopy alone (Figure 2G). For such organelles, fluorescence imaging is indispensable when used in conjunction with ExAPC microscopy. Moreover, we demonstrated the capability of ExAPC microscopy to visualize intracellular structures across various live cell lines, highlighting its broad applicability and potential utility in diverse biological contexts (Figure S1).

**Figure 2.**
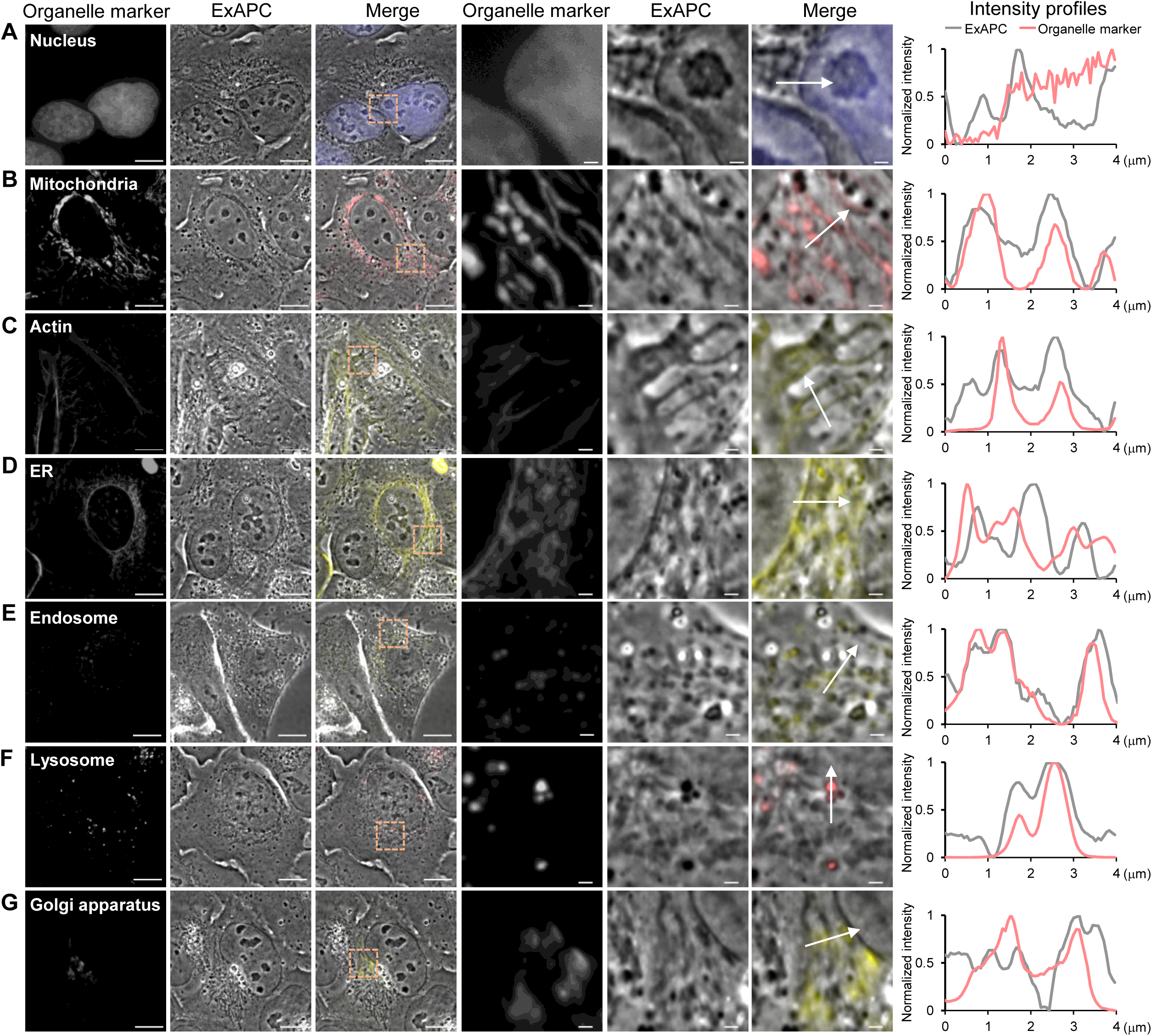
Visualization of cellular organization utilizing ExAPC microscopy. Comparison of organelle visualization and intensity profiles using ExAPC and epifluorescence microscopy with the indicated organelle markers. (A) Nucleus: Hoechst 33342. (B) Mitochondria: MitoTracker Red CMXRos. (C) Actin: Lifeact-EYFP. (D) Endoplasmic reticulum (ER): EYFP-Cb5. (E) Endosome: ECGreen. (F) Lysosomes: LysoTracker Red DND-99. (G) Golgi apparatus: EYFP-sGOLGB1. The intensities obtained by ExAPC microscopy are shown as absolute values after subtracting 1 from the normalized value. Arrows indicate traces of the line scan. Scale bar: 10 μm, scale bar in inset: 1 μm.

Reliable information on the dynamics of cellular organization offers insights into the mechanisms driving various cellular behaviors. ExAPC microscopy utilizes an apodization method to effectively reduce halo artifacts (Figure 1B). As a result, ExAPC microscopy allowed us to capture intricate details of changes in cellular organization during the cell division of HeLa cells and human induced pluripotent stem cells (hiPSCs) (Figure 3A and Videos 2, 3). However, complete removal of halos was not possible in cases where cells underwent behaviors such as rounding or thickening. Specifically, during apoptotic cell structural changes (Figure 3B and Video 4) and the engulfment process during entosis (Figure 3C and Video 5), halo suppression was insufficient. Nevertheless, we captured detailed cellular structures in regions unaffected by halos (Figure 3B, C). Additionally, the ExAPC microscope enabled stable long-term imaging of these cellular behaviors (Figure 3). These findings underscore both the exceptional capabilities and the technical limitations of ExAPC microscopy.

**Figure 3.**
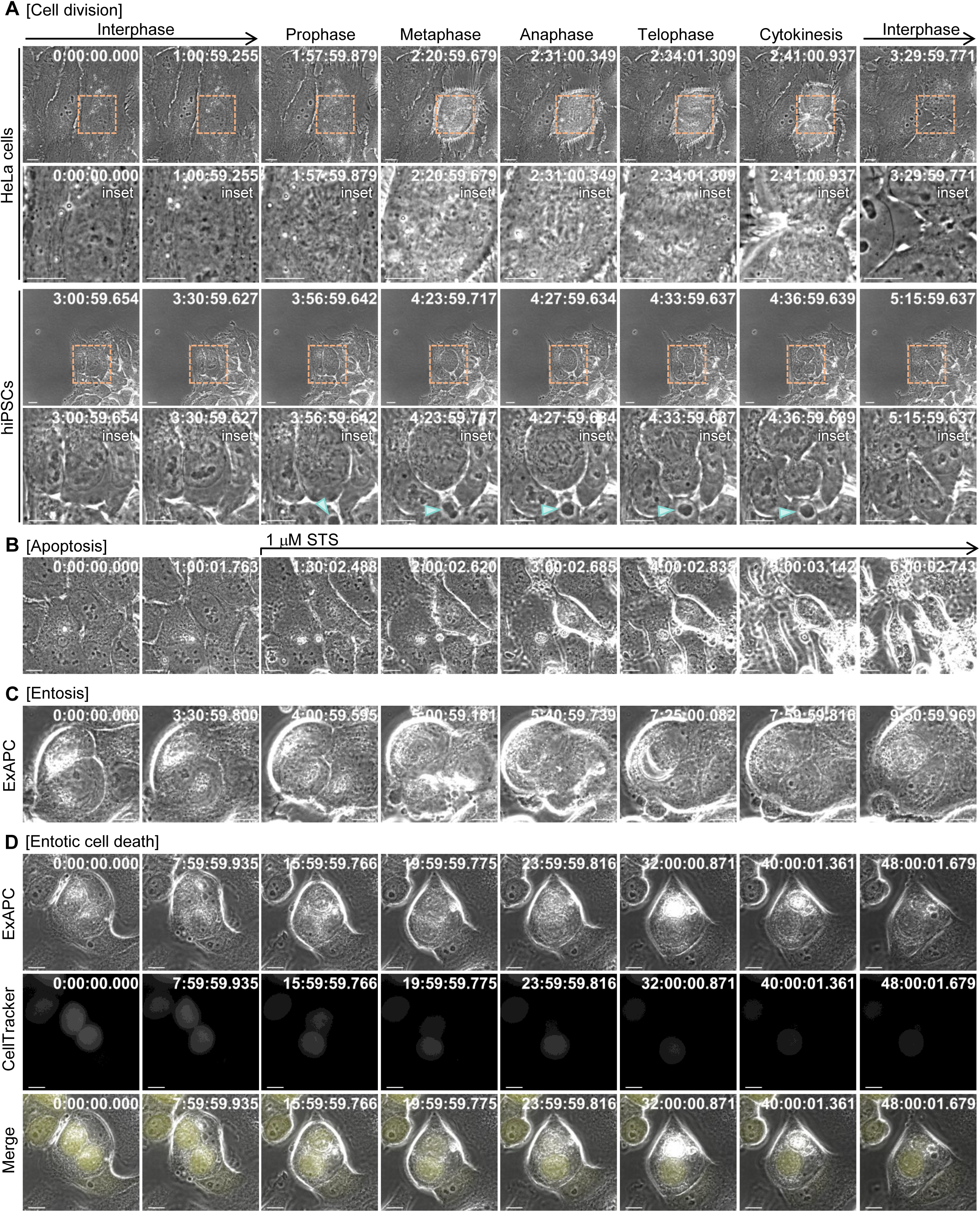
Visualization of diverse cellular behaviors using ExAPC microscopy. Representative images of the indicated cellular behaviors acquired by ExAPC microscopy are shown. (A) Cell division of HeLa cells and human induced pluripotent stem cells (hiPSCs). Acquisition rate: 1 min/frame. Arrowhead: Biomolecular condensate-like structure; see Figure 4. (B) HeLa cells were treated with 1 μM staurosporine (STS) at frame 15. Acquisition rate: 5 min/frame. (C) Entosis of ZR75-1 cells. Acquisition rate: 1 min/frame. (D) ZR75-1 cells stained with CellTracker Green CMFDA and unstained ZR75-1 cells were cocultured. Acquisition rate: 1 min/frame. Scale bar: 10 μm. Time: h:mm:ss.ms.

### ExAPC microscopy-based analysis of biomolecular condensates and their analogous structures

Fluorescence microscopy delineates cellular organization by utilizing intracellular structure-specific fluorescent markers. In contrast, ExAPC microscopy converts the phase shift induced by light traversing the cell into an amplitude shift, resulting in a variation in image contrast. Unlike fluorescence microscopy, ExAPC microscopy simultaneously visualizes all intracellular structures that cause phase shifts within cells without the need for staining.

Using ExAPC microscopy, we examined live HeLa cells and identified highly dynamic and well-defined spherical structures within the cytoplasm (Figure 4A and Video 6). These structures were present in 16.3% (45 of 276 cells) of HeLa cells. A similar structure was observed in hiPSCs (Video 3). There was a weak correlation between the area and average strength of the structures observed in HeLa cells, suggesting that these structures are composed of biomolecules with varying refractive indices (Figure 4B). Time-lapse imaging revealed that these structures underwent fusion events (Figure 4C) and treatment with 1,6-hexanediol (1,6-HD), an agent known to dissolve liquid-phase-separated biomolecular condensates^20^, resulted in their disappearance (Figure 4D). Furthermore, subjecting HeLa cells to hyperosmotic stress, which is known to induce the formation of biomolecular condensates^21^, resulted in the appearance of similar structures in the cytosol (Figure 4E). Based on these findings, we defined these structures as biomolecular condensate-like structures (BCLs). The BCLs displayed dynamic changes in their perimeter (Figure 4F), and localization (Figure 4G) compared to the nucleolus, the major biomolecular condensate in the nucleus^22^. Thus, the mechanisms regulating the structural stability and localization of BCLs may differ from those governing the nucleolus.

**Figure 4.**
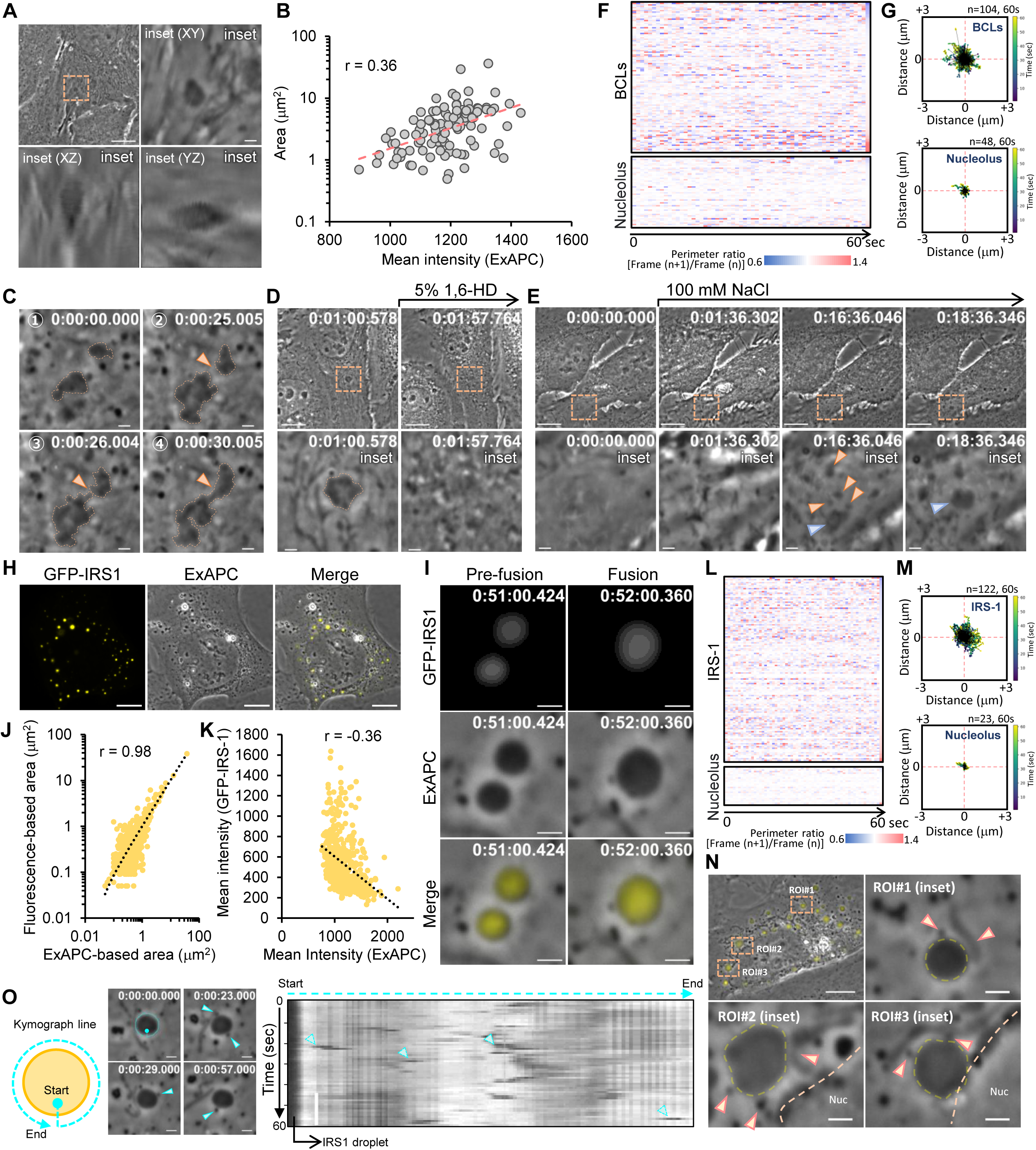
Visualization of BCLs and IRS-1 droplets via ExAPC microscopy. (A) Z-stack imaging of BCL in HeLa cells. Z-Step: 0.097 μm. Scale bar: 10 μm; scale bar in inset: 1 μm. (B) Relationship between the area and average intensity of BCLs observed in HeLa cells. n = 104 from 45 cells. (C) Time-lapse imaging showing the fusion of the two BCLs in HeLa cells. Acquisition rate: 1 sec/frame. Scale bar: 1 μm. (D) BCL-positive HeLa cells were treated with 5% 1,6-HD at Frame 61. Acquisition rate: 1 sec/frame. Scale bar: 10 μm; scale bar in inset: 1 μm. (E) HeLa cells were treated with 100 mM NaCl at Frame 61. Acquisition rate: 1 sec/frame. Scale bar: 10 μm; scale bar in the inset: 1 μm. The BCLs indicated by orange arrowheads were fused with the BCL indicated by the blue arrowhead. (F) Heatmaps of the perimeter ratios of BCLs and nucleoli in HeLa cells. Each row represents a single BCL (n = 104) or the nucleolus (n = 48). (G) Random motions of BCLs and the nucleolus in HeLa cells. The plots show BCLs and nucleoli over 60 s, starting at the center. The analyzed BCLs and nucleolus are the same as that in Figure 4F. (H) Representative images of U-2 OS cells expressing GFP-IRS-1. Scale bar: 10 μm. (I) Time-lapse imaging showing the fusion of two IRS-1 droplets in U-2 OS cells expressing GFP-IRS-1. Acquisition rate: 1 min/frame. Scale bar: 1 μm. (J and K) Relationships between the area of IRS-1 droplets measured from ExAPC and epifluorescence images (J) and the mean intensity of IRS-1 droplets measured from ExAPC and epifluorescence images (K). n = 545 from 15 U-2 OS cells expressing GFP-IRS-1. (L) Heatmaps of the perimeter ratio of IRS-1 droplets to nucleoli in 9 GFP-IRS-1-expressing U-2 OS cells. Each row represents a single IRS-1 droplet (n = 122) or a nucleolus (n = 23). (M) Random motion of IRS-1 droplets and nucleoli in U-2 OS cells expressing GFP-IRS-1. The plots show IRS-1 droplets and nucleoli over 60 s, starting at the center. The analyzed BCLs and nucleolus are the same as that in Figure 4L. (N) Interaction of IRS-1 droplets with various organelles in GFP-IRS-1-expressing U-2 OS cells. Scale bar: 10 μm; scale bar in the inset: 1 μm. (O) Kymograph showing the interaction between the IRS-1 droplets and ELLs (arrowhead). Scale bar: 1 μm. Time: h:mm:ss.ms.

To further validate the capability of ExAPC microscopy in observing biomolecular condensates, we focused on insulin receptor substrate 1 (IRS-1). IRS-1 is a key protein in the regulation of insulin/insulin-like growth factor (IGF) signaling^23–25^, and the phase separation of IRS-1 promotes the formation of the insulin/IGF signalosome, also known as the IRS-1 droplet^26^. Following the exogenous expression of green fluorescent protein (GFP)-fused IRS-1 in U-2 OS cells, we observed the formation of IRS-1 droplets^26^ using both epifluorescence and ExAPC imaging (Figure 4H and Figure S2A). Additionally, ExAPC microscopy revealed the fusion of IRS-1 droplets at various intracellular locations (Figure 4I and Video 7). Notably, 23.5% (167 of 712) of IRS-1 droplets were detectable via epifluorescence imaging, whereas no corresponding structures were visible in the ExAPC images (Figure S2B). Although the IRS-1 droplets did not differ significantly in size between those visible and those invisible to ExAPC microscopy, the fluorescence intensity of droplets not visible in ExAPC imaging was significantly lower (Figure S2C, D). This suggests that the quantity of IRS-1 within IRS-1 droplets may influence their refractive index. Nevertheless, determining the visibility of IRS-1 droplets via ExAPC microscopy based solely on fluorescence intensity proved challenging (Figure S2E).

We then focused on IRS-1 droplets observable by both fluorescence and ExAPC microscopy. The size of IRS-1 droplets was determined to be 0.76 ± 1.90 μm² by epifluorescence imaging and 0.77 ± 1.82 μm² by ExAPC imaging, showing a strong correlation between the two methods (Figure 4J). However, the intensity of IRS-1 droplets in epifluorescence imaging weakly correlated with that observed by ExAPC microscopy (Figure 4K). These results suggest that variations in the quantity and quality of molecules constituting the IRS-1 droplet may influence the refractive index. Time-lapse imaging revealed dynamic changes in both the perimeter and localization of IRS-1 droplets, similar to those seen in BCLs (Figure 4L, M). Furthermore, we used the multiplexing capabilities of ExAPC microscopy to investigate the interactions between IRS-1 droplets and organelles. Consistent with previous findings^27^, we observed interactions between IRS-1 droplets and nuclei, mitochondria, and ELLs (Figure 4N). Notably, interactions with ELLs occurred at high frequencies, with some lasting less than 1 second (Figure 4O). The interactions between IRS-1 droplets and organelles, occurring at various locations and over differing durations, may be critical for optimizing IRS-1-mediated signal transduction.

### Dissection of lipid droplet formation using ExAPC microscopy

Lipid droplets (LDs) are increasingly recognized as intracellular organelles with diverse functions beyond energy storage, attracting interest in understanding their biosynthetic pathways, physiological roles, and involvement in disease processes. However, the mechanisms underlying LD biogenesis and function, as well as their associations with human diseases, remain poorly understood^28^. Although various reagents exist for labeling LDs, LD formation requires extended time, highlighting the need for label-free observation methods to circumvent phototoxicity and photobleaching limitations.

We acquired time-lapse images of HeLa cells treated with 200 µM oleic acid to observe LD biosynthesis in detail using ExAPC microscopy (Figure S3A). Among the structures observed, those stained with LipiBlue, which enhances fluorescence in hydrophobic environments^29^, were identified as LDs (Figure S3B). Consequently, ExAPC microscopy allowed us to track the entire trajectory of LDs from initial appearance to subsequent growth and expansion (Figure 5A, Video 8). When the LD diameter was less than 450.4 ± 70.9 nm, it was observed as a white or black dot-like structure (Figure S3C, D). In contrast, once the LDs exceeded this diameter, they shifted from dot-like to ring-like structures (Figure S3E, F). Notably, the appearance of ring-like LDs varied depending on focal position (Figure S3F). We defined phase 0 as the period when no LDs were observed, followed by phases 1 to 3, which corresponded to LD diameters (Figure 5A).

**Figure 5.**
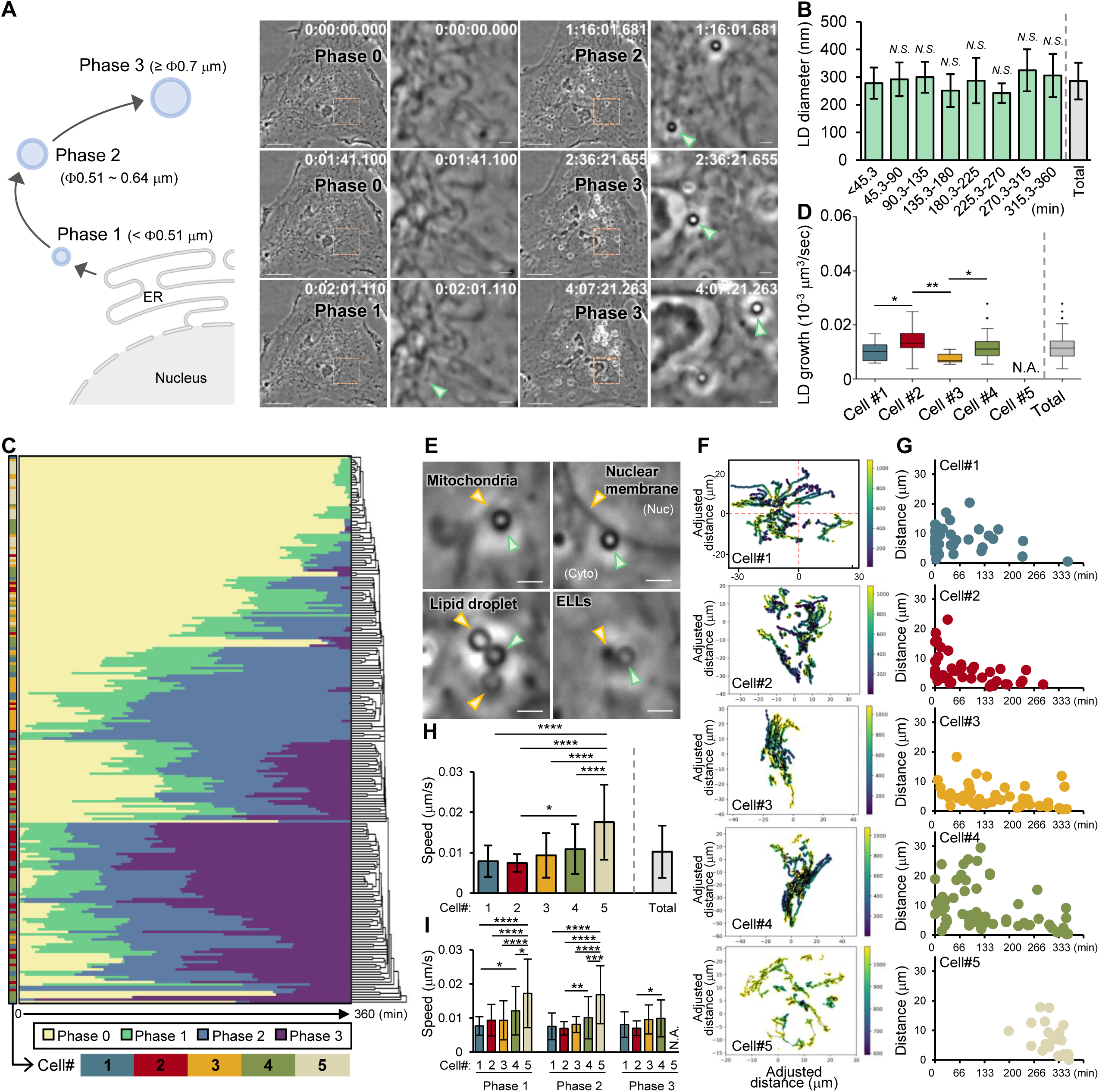
Visualization of LD dynamics using ExAPC microscopy. (A) [left] Growth process of LDs. [right] Time-lapse imaging was started 20 min after adding 200 μM oleic acid to the HeLa cells. See Figure S3A. The cells shown here are the same as in Figure S3B. Green arrowhead: LD being tracked. Scale bar: 10 μm; scale bar in inset: 1 μm. (B) Diameters of LDs recognized as Phase 1 at each time point after image acquisition began (frame 1: 0 min). Acquisition rate: 20 sec/frame. means ± SD. Number of LDs at each period from left to right: 51, 21, 29, 15, 10, 14, 12, 24. One-way ANOVA for multiple comparisons between each period. N.S., no significant difference. Total: all analyzed LDs (n = 176). (C) Heatmap of phase transitions for each LD (n = 218). The cells analyzed (Cell#1-5) and in each phase (Phase 0-3) are color-coded according to the legend below. The dendrogram shows a hierarchical clustering based on the time at which the phase transition occurred for each LD. (D) LD growth rate based on the Phase 2-3 transition time. means ± SD. The numbers of LDs analyzed were as follows: Cell#1: n = 23; Cell#2: n = 35; Cell#3: n = 8; Cell#4: n = 35; Cell#5: n = 0 (N.A.); and Total: n = 101. One-way ANOVA for multiple comparisons was performed between each cell. (E) Interaction of LDs with various organelles in HeLa cells. Green arrowhead: LD. Yellow arrowhead: organelle interacting with the LD. Scale bar: 1 μm. (F) LD dynamics. The coordinates of LDs are corrected for the coordinates of nucleoli in the same cell. The time at which the coordinates of each LD were obtained after the addition of oleic acid is color coded according to the legend on the right. The number of LDs is plotted as follows: Cell#1: n = 36; Cell#2: n = 44; Cell#3: n = 44; Cell#4: n = 65; and Cell#5: n = 28. (G) Euclidean distance between the coordinates at which each LD was first identified and the coordinates at the end of time-lapse imaging. The number of plotted LDs is the same as that in Figure 5F. (H) LD migration velocity in each cell (Cell #1-5) and in all cells (total). Means ± standard deviation. The analyzed LDs is the same as that in Figure 5F. One-way ANOVA for multiple comparisons was performed between each cell. \**P* < 0.05, \*\*\*\**P* < 0.0001. (I) LD migration velocity in each cell (Cell#1-5) in each phase. Means ± standard deviation. The analyzed LDs is the same as that in Figure 5F. The numbers of LDs analyzed were as follows: [Phase 1] Cell#1: n = 33, Cell#2: n = 42, Cell#3: n = 42, Cell#4: n = 43, Cell#5: n = 28, [Phase 2] Cell#1: n = 36, Cell#2: n = 44, Cell#3: n = 38, Cell#4: n = 58, Cell#5: n = 10, [Phase 3] Cell#1: n = 25, Cell#2: n = 37, Cell#3: n = 8, Cell#4: n = 53, Cell#5: n = 0. One-way ANOVA for multiple comparisons was performed between each cell in each phase. \**P* < 0.05, \*\**P* < 0.01, \*\*\**P* < 0.001, \*\*\*\**P* < 0.0001.

When the 218 LDs that were positive for Lipi-Blue in the final frame were analyzed, 176 LDs appeared as phase 1 LDs during time-lapse imaging, with a mean diameter of 285.6 ± 66.2 nm (Figure 5B). This diameter is similar to that of pre-LDs (257.6 ± 31 nm), an ER microdomain containing a stable core of neutral lipids in early LD biogenesis^30^. The diameter of phase 1 LDs remained constant over time following oleic acid addition (Figure 5B). The time required for LDs to transition to phases 1, 2, and 3 was 143.3 ± 116.4 min (n = 176), 161.9 ± 92.1 min (n = 162), and 239.6 ± 76.4 min (n = 123), respectively, after imaging began (Figure S3G). The mean transition times varied significantly among cells (Figure S3G). LD growth was heterogeneous across cells following oleic acid addition (Figure 5C). The time for LDs to grow from phase 2 to phase 3 was 110.3 ± 44.0 min (n = 101), yielding an average growth rate of 0.012 ± 0.004 × 10⁻³ µm³/sec (Figure 5D). LD growth rates also varied significantly among cells (Figure 5D). Notably, no LD size changes due to fusion or division were observed during the imaging period.

LDs interact frequently with organelles, serving as lipid sources for energy production and signal transduction^31^. Using ExAPC microscopy, we observed interactions between LDs and various organelles, including mitochondria and the nucleus, without the need for labels (Figure 5E). Additionally, tracking LD dynamics revealed that LDs do not remain stationary but instead migrate to specific subcellular compartments (Figure 5F, G). The migration velocity of LDs was 0.01 ± 0.006 μm/sec, with statistically significant variation between cells (Figure 5H, I).

Taken together, these observations highlight the complex behavior and dynamic nature of LDs within the cellular environment, including their interactions with other organelles and spatial redistribution across different cellular compartments.

### Mitochondrial dynamics analyzed using ExAPC microscopy

Mitochondria are dynamic organelles that undergo tightly regulated fission and fusion events^32^. Disruption of these regulatory mechanisms has been linked to various diseases, including cancer, cardiovascular diseases, and neurodegenerative disorders^33^. Therefore, observing mitochondrial morphology is crucial for clarifying pathogenetic mechanisms. However, assessing mitochondrial morphology through fluorescence imaging requires mitochondria-specific probes and precise imaging parameters, such as light intensity and exposure time^34^. In contrast, mitochondrial morphology can be visualized unambiguously and in a label-free manner using ExAPC microscopy. Compared to fluorescence imaging, ExAPC microscopy is more convenient and enables more detailed longitudinal observations.

We utilized ExAPC microscopy to investigate spontaneous mitochondrial fission and fusion processes without pharmacological or genetic induction. As a result, we accurately delineated the locations of fission in mitochondria of various shapes and lengths, similar to observations made using fluorescence imaging^35^ (Figure 6A). Mitochondrial fission is driven by the cooperative action of the ER and dynamin-related protein 1 (DRP1) in constricting mitochondrial cleavage sites^36,37^. This constriction was detectable via ExAPC microscopy (Figure 6B and Video 9). The fission sites were observed at various locations along the mitochondria, with the distribution displaying a bimodal pattern (Figure 6C). This observation supports the existence of two distinct mitochondrial fission mechanisms, as previously reported^35^. We further examined whether mitochondrial length influenced fission site location but found no significant correlation between mitochondrial length and the fission site (Figure 6D).

**Figure 6.**
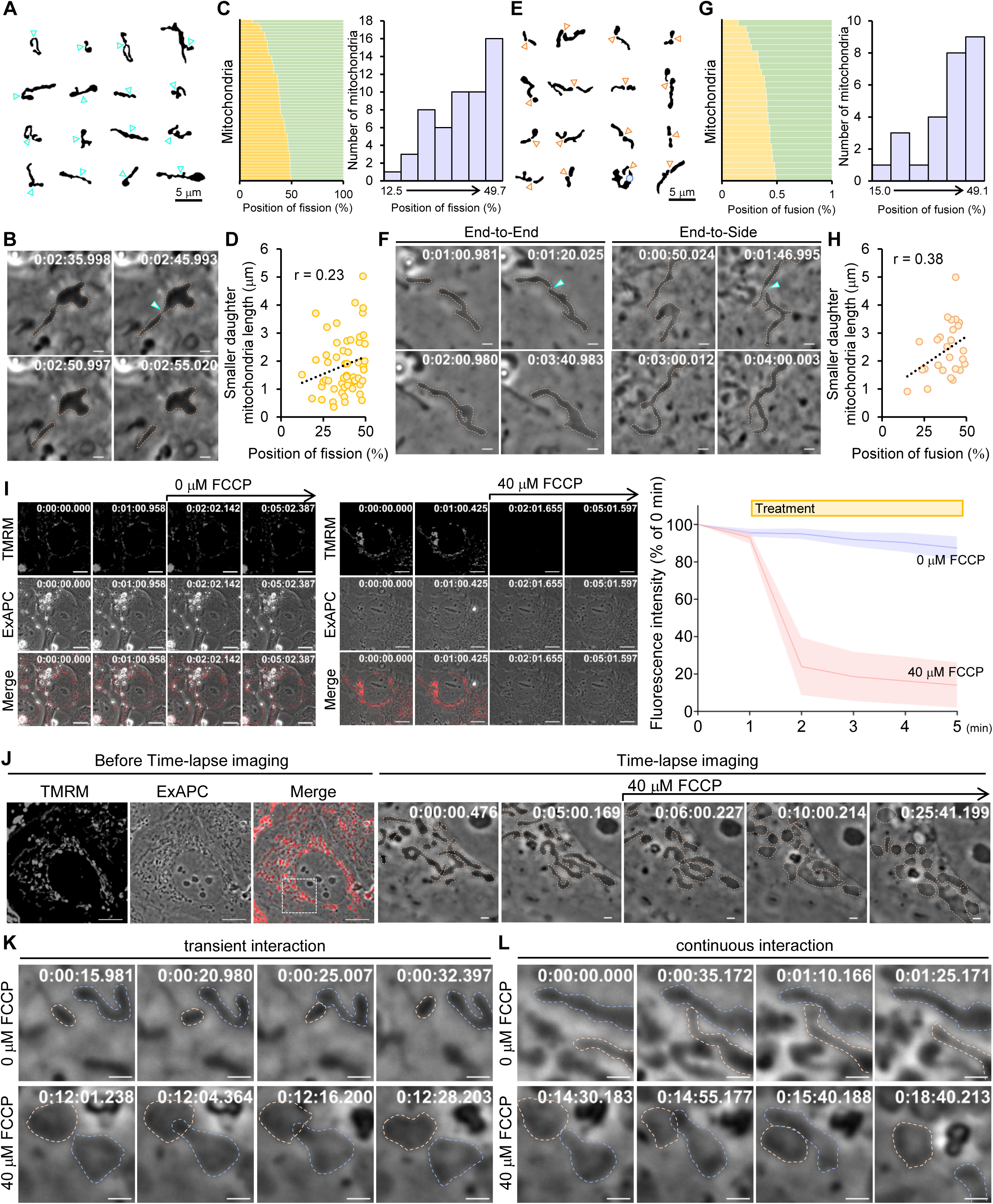
Visualization of mitochondrial dynamics using ExAPC microscopy. (A) Gallery image of mitochondria from a binarized ExAPC microscopy Video one frame before division. (B) Time-lapse imaging of mitochondrial fission in Hep3B cells. (C) [Left] Position of mitochondrial fission relative to total mitochondrial length (n = 54 fissions pooled from multiple datasets). [Right] Histogram of fission positions relative to the total mitochondrial length in the left figure. Width: 5.32. (D) Correlation between shorter daughter mitochondria length and the position of mitochondrial fission relative to total mitochondrial length. The analysis of mitochondria was identical to that in Figure 5C. (E) Gallery image of mitochondria from a binarized ExAPC microscopy Video one frame before fusion. (F) Time-lapse imaging of mitochondrial fusion in Hep3B cells. (G) [Left] Position of mitochondrial fusion relative to total mitochondrial length (n = 26 fusions pooled from multiple datasets). [Right] Histogram of fusion positions relative to the total mitochondrial length in the left figure. Width: 5.69. (H) Correlation between shorter daughter mitochondria length and the position of mitochondrial fusion relative to total mitochondrial length. The analysis of mitochondria was identical to that in Figure 5G. (I) Mitochondrial membrane potential in Hep3B cells. Representative images at each time point are shown. All the data are presented as the mean ± standard deviation. 0 μM FCCP: n = 14 cells. 40 μM FCCP: n = 49 cells. (J) FCCP treatment-induced changes in mitochondrial morphology in Hep3B cells. FCCP was added after image acquisition at frame 301 (1 frame/sec). Scale bar: (Before Time-lapse imaging): 10 μm, (Time-lapse imaging): 1 μm. (K and L) Inter-mitochondrial communication in the presence or absence of FCCP in Hep3B cells. Scale bar: 1 μm.

The mitochondrial fusion process was similarly observed. Mitochondrial fusion consists of a two-step process in which mitofusins (MFNs) mediate outer mitochondrial membrane fusion, and optic atrophy-1 (OPA1) regulates inner mitochondrial membrane fusion in mammalian cells^38^. Typically, mitochondrial fusion occurs at the ends of both mitochondria, although end-to-side fusion can occur in rare instances^39^ (Figure 6E, F and Video 10, 11). Similar to fission, a gradual bimodal distribution was observed for the relative lengths of the two fused mitochondria (Figure 6G). However, the relative length of the smaller mitochondria was greater than 30% in 80.7% of fusion events, indicating that mitochondria of similar length tended to fuse (Figure 6G). As with fission, we found no substantial correlation between mitochondrial length and fusion site (Figure 6H). These findings suggest that the ratio of mitochondrial lengths should be considered when examining the mechanisms governing mitochondrial fission and fusion.

Exposure to an oxidative phosphorylation uncoupler, such as carbonylcyanide-p-trifluoromethoxyphenylhydrazone (FCCP), leads to a reduction in mitochondrial membrane potential (ΔΨm) and induces reactive oxygen species (ROS) stress within the cell, causing mitochondria to transform from elongated string-like structures into small globular space^40^. However, the characteristics of mitochondrial morphology following ΔΨm loss remain debated. To explore these characteristics, we employed ExAPC microscopy to visualize the rapid changes in mitochondrial morphology upon FCCP treatment. Specifically, treatment with 40 µM FCCP significantly reduced ΔΨm (Figure 6I and Figure S4A), and the morphology of individual mitochondria in Hep3B cells transitioned from a string-like to a globular form within seconds (Figure 6J and Video 12). In contrast, only a subset of mitochondria in U-2 OS cells adopted a spherical morphology (Figure S4B and Video 13). Furthermore, these morphological changes varied between individual mitochondria in both Hep3B and U-2 OS cells (Figure 6J and Figure S4B). These results suggest heterogeneity in mitochondrial responses to FCCP.

Mitochondria are known for their ability to communicate through physical contact^41^. We observed this inter-mitochondrial communication using ExAPC microscopy, both under normal culture conditions and in conditions where ΔΨm was disrupted by FCCP treatment (Figure 6K, L). These findings suggest that inter-mitochondrial communication can occur independently of ΔΨm.

## Discussion

In this study, we demonstrate that ExAPC microscopy is a valuable tool for examining cellular organization in diverse cell types, such as human cancer cell lines, primary cultured cells, and iPS cells. With its high spatiotemporal resolution and multiplexing capabilities, this label-free imaging modality facilitates simultaneous visualization of various intracellular structures and allows detailed analysis of morphological changes in cellular organization across a range of physiological and pathological contexts. Despite recent advances in photostable fluorescent proteins^42,43^, label-free imaging provides distinct advantages, particularly enabling prolonged observations of living cells with minimal concerns about phototoxicity and photobleaching.

We obtained quantitative data on biomolecular condensates, lipid droplets, and mitochondria using ExAPC microscopy, and comparable results can be achieved using conventional fluorescence imaging, except for BCLs. Therefore, ExAPC microscopy offers a significant advantage in its label-free imaging capability, enhancing experimental flexibility. For instance, in label-free imaging, there is no need to account for the potential effects of staining on the cells, allowing for subsequent quantification of gene or protein expression levels after observation using ExAPC microscopy. Moreover, the reduced risk of phototoxicity and photobleaching in ExAPC microscopy enables extended real-time observation of cellular dynamics, which is a major benefit. Selecting the optimal microscopic technique for specific experimental goals is critical for advancing science research. ExAPC microscopy is well suited for experiments requiring high temporal resolution or the observation of thin-cell specimens, while ODT is more effective for thicker samples or when temporal resolution is less critical. Fluorescence imaging remains indispensable for studying organelles, such as the Golgi apparatus, or specific protein activities that cannot be observed using ExAPC microscopy or ODT. Combining these techniques offers new insights into cellular structure and function, potentially driving significant advances in life science research.

Biomolecular condensates are membrane-free intracellular compartments that orchestrate diverse biochemical reactions with spatial and temporal precision^44,45^. However, critical uncertainties remain regarding the processes that govern their formation and the resulting modifications to biomolecule functionality. Using ExAPC microscopy, we observed BCLs for the first time. Further research is needed to elucidate the composition and function of BCLs, but their observation using ExAPC microscopy is expected to enhance our understanding of biomolecular condensates. Additionally, we discovered that the refractive index of IRS-1 droplets varies, suggesting that these droplets consist of distinct molecular components. These findings underscore the limitations of fluorescence imaging, which relies on labeling target proteins, and highlight the advantages of integrating ExAPC microscopy with fluorescence techniques. Over the past decade, researchers have sought to elucidate the role of biomolecular condensates in diseases such as neurodegeneration and cancer^46–48^. It may be crucial to consider the heterogeneity of biomolecular condensates in future research.

Intracellular LDs are conserved organelles present in all eukaryotic cells, storing lipids such as TAGs and cholesterol esters while protecting cells from lipotoxicity by sequestering fatty acids^49,50^. LDs form when TAGs accumulate in the ER membrane and bud into the cytosol, where they grow through fusion or induce de novo TAG synthesis. Fusion involves transferring neutral lipids from smaller to larger LDs via proteins, such as FSP27 and perilipin1, though perilipin1^51^ is mainly found in specific tissues and not in HeLa cells^52,53^. Therefore, in our study, although we observed adjacent LDs over a long period of time, we did not detect any fusion events between them.

Early LDs are known for their short lifespan and small size, which often pose challenges to their visualization^50^. Previous studies using electron microscopy have shown that the diameter of nascent LDs varies, with studies on yeast indicating a diameter of 30-60 nm^54^, and studies on mammalian cell lines a diameter of approximately 170 nm^55^. Notably, when ExAPC microscopy was employed to investigate the nascent stages of LD biogenesis, the diameter of newly formed LDs was measured at 285.6 ± 66.2 nm. Observing these early LDs requires high-magnification objectives, and the ExAPC method is well suited for this purpose owing to the capacity to select appropriate lenses. Furthermore, in our study, the growth rate of LDs was calculated to be 0.012 ± 0.004 × 10^-3^μm^3^/sec, a growth rate closer to that observed in COS-1 cells (0.043 × 10^-3^ μm^3^/sec)^30^ than to the rate observed in Drosophila (0.159 × 10^-3^ μm^3^/sec)^56^. Thus, label-free imaging using the ExAPC microscope is a powerful tool for quantitatively analyzing the biogenesis of LDs and their spatial relationships with other intracellular structures. By applying this technology, we can elucidate previously unresolved aspects of LD function and roles, offering the potential to gain new insights into the contribution of LD to diseases such as diabetes and metabolic-associated fatty liver disease (MAFLD) in various tissues, including adipose tissue and the liver, where LD is said to be important^57,58^.

Mitochondria are multifunctional organelles that undergo dynamic morphological changes through repeated fission and fusion processes^59^. The mitochondria exhibit a plethora of distinct morphologies within different cells, ranging from reticulation throughout the cell to fragmentation into discrete pieces^60^. A series of distinctive morphological alterations in mitochondria have been described during processes such as stem cell differentiation and multifactorial diseases, such as cancer, cardiovascular diseases, and neurodegenerative diseases^5,61,62^. These observations highlight the significance of mitochondrial morphology in various physiological and pathological processes. Understanding the regulatory mechanisms governing mitochondrial fission and fusion is crucial, as these processes determine mitochondrial morphology. In this study, specific patterns were observed regarding the sites of mitochondrial fission and the lengths of two mitochondria undergoing fusion. While the underlying mechanisms governing these regularities remain elusive, mitochondria may possess an overarching ability to perceive their own length and that of neighboring mitochondria, potentially guiding these processes. We also found that ΔΨm loss leads to distinct morphological changes, with Hep3B mitochondria becoming spherical, while U-2 OS mitochondria displayed partial spherical morphology. This discrepancy may stem from differences in mitochondrial lipid composition. Despite the universality of mitochondrial function, questions remain about how differences in mitochondrial morphology and responsiveness to stimuli affect function. In our study, individual mitochondria responded differently to FCCP treatment, suggesting that heterogeneity in mitochondrial responsiveness may play a role in regulating overall mitochondrial function. This variability in cellular responses has been reported to enable more precise control of tissue responses^63^, and similar regulatory systems may exist within mitochondria. Further investigations are required to address these questions. We believe that ExAPC microscopy data will stimulate further exploration of these biological questions and contribute to a deeper understanding of mitochondrial biology.

The choice of microscopy modality depends on the sample characteristics and the need for temporal versus spatial resolution. To better understand cellular behavior, it is essential to observe cells using multiple modalities and comprehensively interpret the data. Recent advances in single-cell omics technologies have enabled real-time omics analysis^64–66^. Integrating these technologies with label-free imaging techniques like ExAPC microscopy, which offers multiplexing capabilities, represents a key challenge for the future of imaging technology.

## Materials and Methods

### Plasmid construction

EYFP-Cb5 was produced by subcloning the sequence coding aa 100–134 of rat Cytochrome b5 (UniprotKB-P00173) into an EYFP vector at the C-terminus between the EcoR I and Sal I sites. EYFP-sGOLGB1 was produced by subcloning the sequence encoding aa 3131–3259 of human GOLGB1 (UniProtKB-Q14789) into an EYFP vector at the C-terminus between the EcoR I and Sal I sites. Lifeact-EYFP was kindly provided by Dr. Hideki Nakamura (Kyoto University). GFP-rIRS-1 was produced by subcloning the sequence coding aa 4–1235 of rat IRS-1 (UniProtKB-P35570) into an EGFP vector at the C-terminus between Hind III and the BamH I site.

### Cell culture and transfection

Human lung carcinoma A549 cells (CCL-185), human cervical adenocarcinoma HeLa cells (CCL-2), human colon carcinoma HCT116 cells (CCL-247), human hepatocellular carcinoma Hep 3B cells (HB-8064), mouse fibroblast NIH/3T3 cells (CRL-1658), human breast adenocarcinoma MDA-MD-231 cells (CRM-HTB-26), human osteosarcoma U2-OS cells (HTB-96), and human breast carcinoma ZR75-1-1 (CRL-1500) cells were purchased from the American Type Culture Collection. These cells, except ZR75-1-1, were cultured in Dulbecco’s modified Eagle’s medium (DMEM; Thermo Fisher Scientific, 11965118) supplemented with 10% fetal bovine serum (FBS; Thermo Fisher Scientific, 10270-106) and 1% Zell Shield (Minerva Biolabs GmbH, 13-0050) at 37°C in 5% CO_2_. ZR75-1-1 cells were cultured in RPMI 1640 medium (Thermo Fisher Scientific, 11875093) supplemented with 10% FBS and 1% Zell Shield at 37°C in 5% CO_2_. For transient transfection, 2 × 10^5^ cells were plated on a 35 mm imaging dish with a polymer coverslip bottom (ibidi, 80136) and incubated for 6 h at 37°C in 5% CO_2_. Following incubation, the cells were transfected with the plasmid using FuGENE HD (Promega, E2311). The indicated experiments were carried out 12–18 h after transfection. The TKT3V1-7 iPSC clone was generated from peripheral blood T cells by Sendai virus reprogramming vectors as previously described^67^. The iPSC line was provided by CiRA with the approval of the Ethical Review Board of CiRA (no. G590). iPSCs were maintained on an iMatrix-511 (Matrixome, Japan) as previously described^68^, but StemFit AK02N (Ajinomoto, Japan) was used instead of AK03.

### Primary astrocyte and neuron culture

Primary culture was performed as previously described^69,70^, with slight modifications. Astrocytes were prepared from the hippocampus on postnatal day 2 and neurons on embryonic day 17.5. Hippocampal extraction was performed as previously described^71^. Cells were isolated using Neural Tissue Dissociation Kits (Miltenyi Biotech, 130-093-231). Astrocytes were then seeded at 5 × 10^5^ cells in 25 mm^2^ flasks (VIOLAMO, 2-8589-01) coated with poly-L-lysine (Merck, P4707), and after 7 days, cells were detached by trypsin treatment and seeded at 3 × 10^5^ cells in 35 mm imaging dishes with polymer coverslip bottoms coated with poly-L-lysine and cultured for 7 days. Neurons were seeded at 5 × 10^5^ cells in 35 mm imaging dishes with a polymer coverslip bottom coated with poly-L-lysine and cultured for 10 days. Astrocytes were cultured in Dulbecco’s modified Eagle’s medium (DMEM) supplemented with 10% fetal bovine serum and 1% penicillin-streptomycin (Nacalai tesque, 09367-34) at 37°C in 5% CO2. Neurons were cultured in a Neurobasal medium (Thermo Fisher Scientific, 21103049) supplemented with 1% B-27 (Thermo Fisher Scientific, 17504044), 1% GlutaMAX (Thermo Fisher Scientific, 35050061), and 1% penicillin-streptomycin at 37°C in 5% CO2.

### Live-cell imaging

For ExAPC microscopy, an apodized phase plate was inserted into a relayed objective’s pupil plane in an inverted Eclipse Ti2-E microscope (Nikon) to obtain images without phase-contrast objectives (the motorized external phase-contrast system, Nikon). Apodized phase contrast images were obtained using an oil-immersion objective (CFI Apochromat TIRF 100XC Oil, 1.49NA, Nikon). To compare the images with conventional phase-contrast images, an oil-immersion phase-contrast objective (CFI Plan Apo DM Lambda 100X, 1.45NA, Nikon) was used. For epifluorescence microscopy images, BFP, EYFP, and mCherry excitation was carried out using an Intensilight mercury-fiber illuminator (Nikon). The data were processed through a BFP-A-Basic filter (Semrock), a CFP-A-Basic-NTE filter (Semrock), a YFP-A-Basic-NTE filter (Semrock), and an mCherry-B-NTE-ZERO filter (Semrock) for BFP, EYFP, and mCherry imaging, respectively. The cells were viewed using a 100× objective mounted on an inverted Eclipse Ti2-E microscope (Nikon) and imaged using a Zyla 4.2 PLUS sCMOS camera (Oxford Instruments). Imaging data were processed using NIS-Elements AR imaging software (Nikon). For ODT, an inverted Eclipse Ti2-E microscope (Nikon) was used, and bright-field images were acquired using a Digital Sight 50M monochrome CMOS camera (Nikon). Bright-field images were obtained using an oil-immersion lens (CFI Plan Apochromat Lambda 100XH, NA = 1.45, Nikon). Imaging data were processed according to previously reported formula^72^. All imaging experiments were completed at 37°C in 5% CO_2_ using an STX stage top incubator (Tokai-Hit). For all live-cell imaging, unless otherwise noted, cells were cultured in phenol red-free DMEM (Thermo Fisher Scientific, 31053028) supplemented with 10% FBS, 4 mM L-glutamine (Thermo Fisher Scientific, 25030081), and 1% penicillin‒streptomycin (Sigma‒Aldrich, P4333). The following representative images obtained by epifluorescence microscopy were processed by Clarify.ai and Denoise.ai using NIS-Elements AR 5.30: mitochondria, ER, Golgi apparatus, and lysosome.

### Organelle staining

A total of 2 × 10^5^ cells were plated on a 35 mm imaging dish with a polymer coverslip bottom and incubated for 12–18 h at 37°C in 5% CO_2_. Following incubation, the cells were cultured with 0.1 μg/ml Hoechst 33342 (Thermo Fisher Scientific, 62249) for 10 min at 37°C in 5% CO_2_ to stain the nuclei. For mitochondrial staining, the cells were cultured with 0.5 μM MitoTracker Red CM-H2Xros (Thermo Fisher Scientific, M7513) for 30 min at 37°C in 5% CO_2_. The endosomal staining was performed using the ECGreen-Endocytosis Detection kit (DOJINDO, E296) in accordance with the provided protocol. For lysosome staining, the cells were cultured with 100 nM LysoTracker Red DND-99 (Thermo Fisher Scientific, L7528) for 30 min at 37°C in 5% CO_2_. The cells were then washed with imaging medium twice and subjected to live-cell imaging.

### Monitoring cell division

For the HeLa cells, 2 × 10^5^ cells were plated on a 35 mm imaging dish with a polymer coverslip bottom and incubated for 24 h at 37°C in 5% CO_2_. For iPSCs, 1.3 × 10^5^ cells were plated on a 35 mm imaging dish with a polymer coverslip bottom at 37°C in 5% CO_2_. After 72 h, live-cell imaging was carried out with the iPSC culture medium.

### Monitoring of apoptosis

HeLa cells (2 × 10^5^ cells) were plated on a 35 mm imaging dish with a polymer coverslip bottom and incubated for 24 h at 37°C in 5% CO_2_. Subsequently, the cells were subjected to live-cell imaging. Then, 1 μM staurosporine (Sigma‒Aldrich, S6942-200UL) was added at the indicated times.

### Monitoring entosis

ZR75-1-1 cells (4 × 10^5^ cells) were seeded in three 6 cm dishes. Forty-eight hours later, 4 µl of 1 mM CellTracker Green CMFDA Dye (Thermo Fisher Scientific, C7025) was added to one dish and incubated for 30 min at 37°C in 5% CO_2_. After incubation, the cells were washed 3 times with 4 ml of PBS. The ratio of stained cells to unstained cells was adjusted to 1:2, and these cells were plated onto 35mm imaging dishes with polymer coverslips (4 × 10^5^ cells/dish) and incubated for 48 h at 37°C in 5% CO_2_. Subsequently, the cells were subjected to live-cell imaging.

### Monitoring BCLs and IRS-1 droplets

HeLa cells (2 × 10^5^ cells) were plated on a 35 mm imaging dish with a polymer coverslip bottom and incubated for 24 h at 37°C in 5% CO_2_. Subsequently, the cells were subjected to live-cell imaging. 1,6-Hexanediol (Sigma‒Aldrich, 240117) and sodium chloride (Nacalai, 31320-05) were added at the indicated times. The perimeter, intensity, and motility of the BCLs and IRS-1 droplets were analyzed by NIS software (Nikon).

### LD tracking and analysis

HeLa cells (2 × 10^5^ cells) were plated on a 35 mm imaging dish with a polymer coverslip bottom and incubated for 24 h at 37°C in 5% CO_2_. The cells were stained with 0.5 μM Lipi-Blue for 30 min according to the manufacturer’s protocol. Oleic acid (200 µM; Sigma‒Aldrich, O1008) was then added to the cells 20 min before live-cell imaging began. LD tracking was performed using NIS software (Nikon). For the classification of LDs from Phase 1 to Phase 3, LDs with diameters ranging from 0.51 to 0.64 μm were classified as Phase 2, while LDs smaller than this range were categorized as Phase 1, and those larger than this range were categorized as Phase 3. In calculating the growth rate of LDs from Phase 2 to Phase 3, only LDs that exhibited a transition from Phase 1 to Phase 3 during time-lapse imaging were included in the calculation. Heatmap cluster analysis was performed by Origin Pro 2023. For plotting the behavior of LDs, the coordinates of the nucleolus in the same cell were subtracted from the coordinates of the LD.

### ΔΨm analysis

The cells were cultured for 30 min in the presence of 100 nM Image-iT TMRM Reagent (TMRM; Thermo Fisher Scientific, I34361) at 37°C in 5% CO_2_. The cells were then washed with imaging medium twice, and TMRE fluorescence in the mitochondria was recorded with NIS software (Nikon). Background fluorescence was measured in areas without mitochondria and subtracted from the fluorescence obtained from mitochondria.

### Statistical analysis

The results are presented as the means ± standard deviation. All the statistical tests were performed using GraphPad Prism (GraphPad Software) or Excel (Microsoft Office). The number of classes in the histogram is determined by the Sturgess formula.

## Acknowledgments

We thank Katsuko Okubo (University of Tsukuba) for her technical assistance and Yasuko Maruyama (University of Tsukuba) for her generous support. We would like to thank Nature Research Editing Service for English language editing. This study was supported in part by a JSPS KAKENHI (22H02296), the Takeda Science Foundation (DGM06005J), JST COI-NEXT (JPMJPF2017), JST CREST (JPMJCR1927), and Cabinet Office, Government of Japan, Cross-ministerial Moonshot Agriculture,Forestry and Fisheries Research and Development Program, “Technologies for Smart Bio-industryand Agriculture”(funding agency: Bio-oriented Technology Research Advancement Institution) (JPJ009237).

## Author contributions

T.Miyamoto. conceived the project. T.Miyamoto. designed the experiments. T. Miyamoto., H.O., T.N., K.K., Y.O., N.O., M.K., L.N.S., M.M., Y.Y., Y.H., Y.Y., SI. T., Y.M., Y.Y., Y.T., M.S., T.M., H.S. conducted experiments. T.Miyamoto., and H.O. wrote the manuscript.

## Figure legends

**Figure S1.**
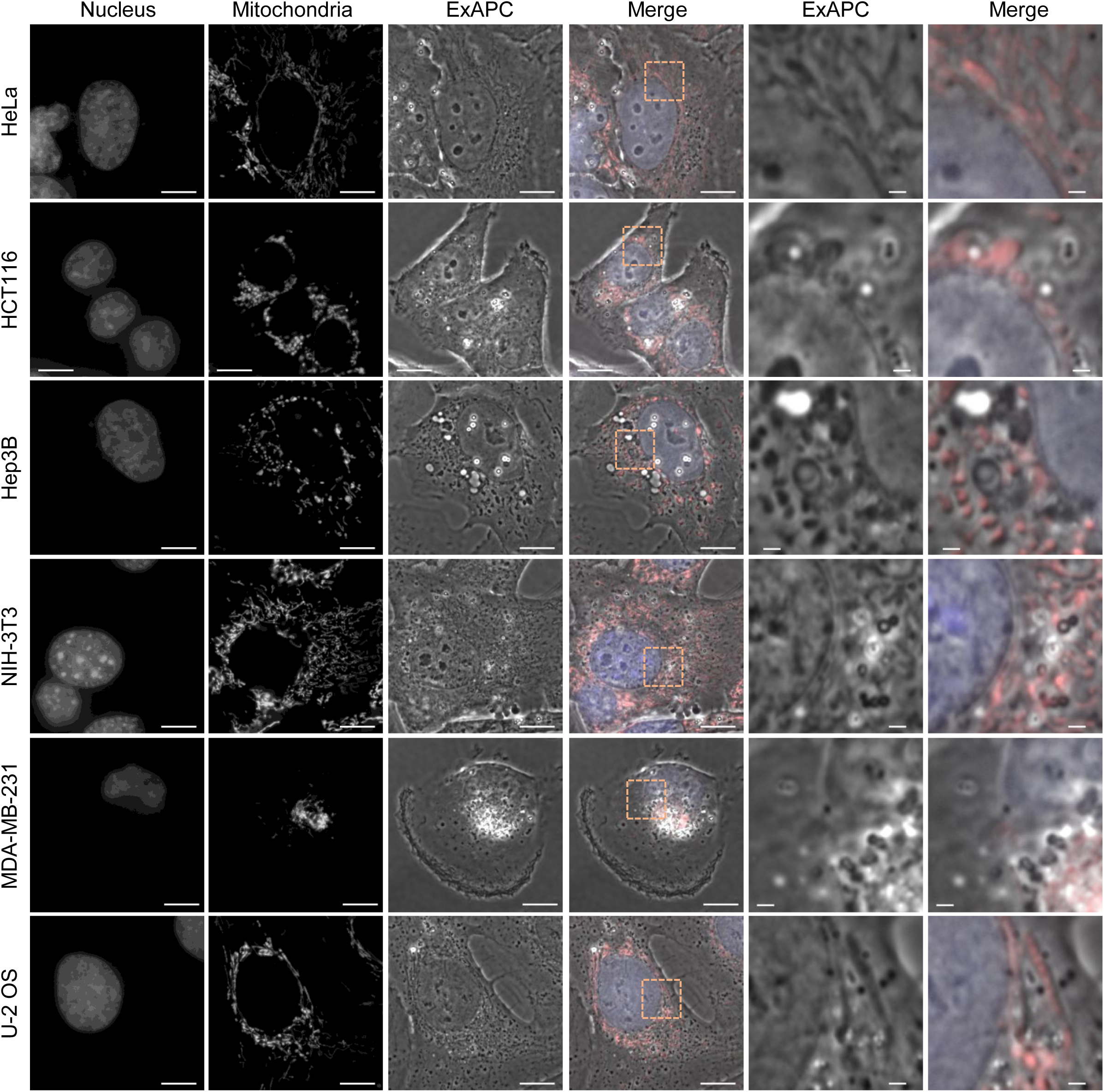
Various cell lines visualized by ExAPC microscopy. Representative images of the indicated cell lines acquired by ExAPC microscopy are shown. Nucleus: Hoechst 33342; mitochondria: MitoTracker Red CMXRos. Scale bar: 10 μm, Scale bar in inset: 1 μm.

**Figure S2.**
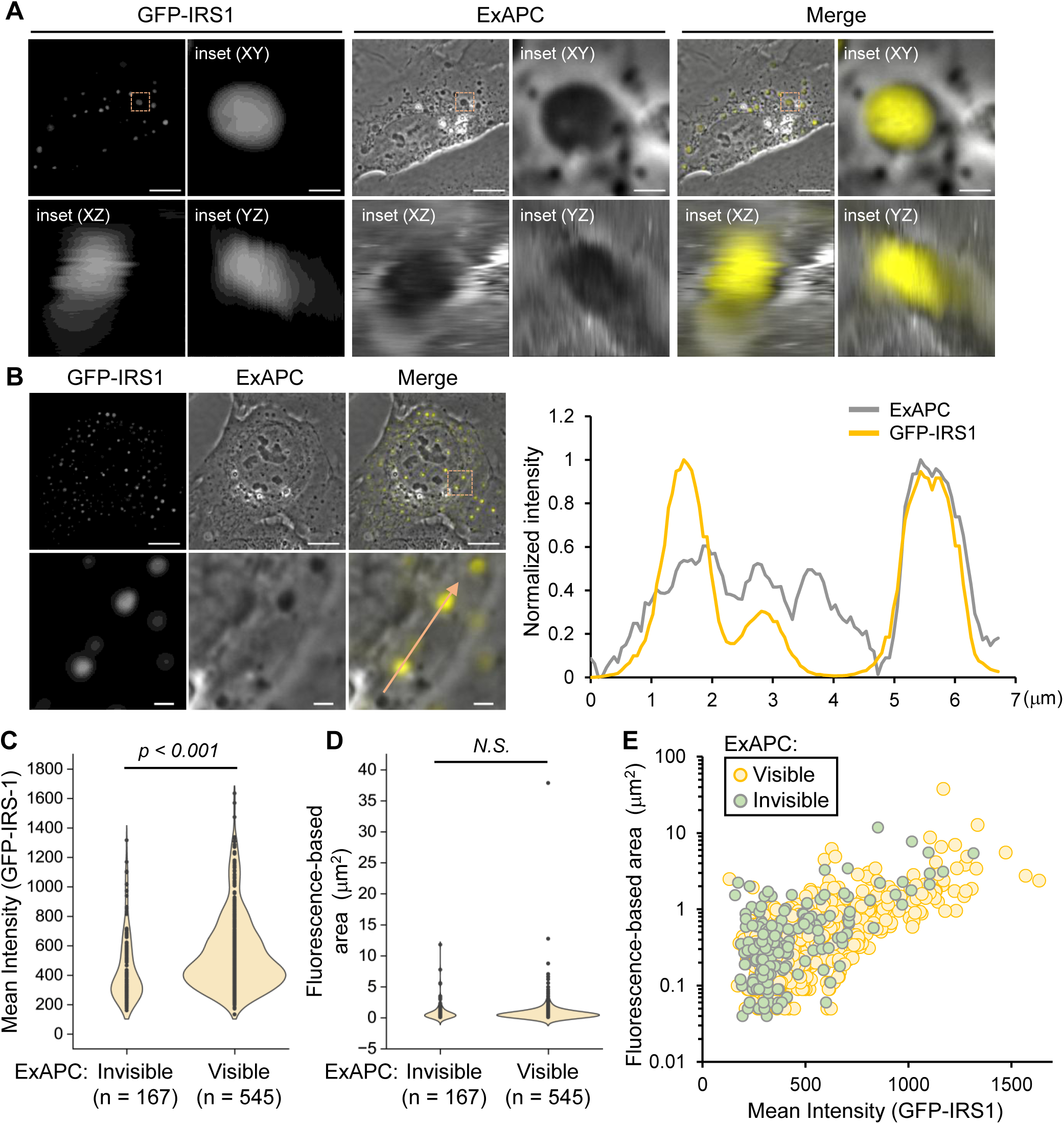
Visibility of IRS-1 droplets via ExAPC microscopy. (A) Z-stack imaging of IRS-1 droplets in U-2 OS cells. GFP-IRS-1 was transiently expressed in U-2 OS cells. Z-Step: 0.048 μm. Scale bar: 10 μm, Scale bar in inset: 1 μm. (B) line-scan profiles of intensity in the merged inset is shown. The intensities obtained by ExAPC microscopy are shown as absolute values after subtracting 1 from the normalized value. Scale bar: 10 μm; scale bar in inset: 1 μm. (C and D) Mean fluorescence intensity (C) and area (D) of IRS-1 droplets that cannot be detected in GFP-IRS-1-expressing U-2 OS cells via ExAPC microscopy. p: Unpaired t-tests. N.S., statistically nonsignificant. (E) Relationship between the IRS-1 droplet area and the mean intensity of IRS-1 in U-2 OS cells expressing GFP-IRS-1, as measured in Figure S2C, D.

**Figure S3.**
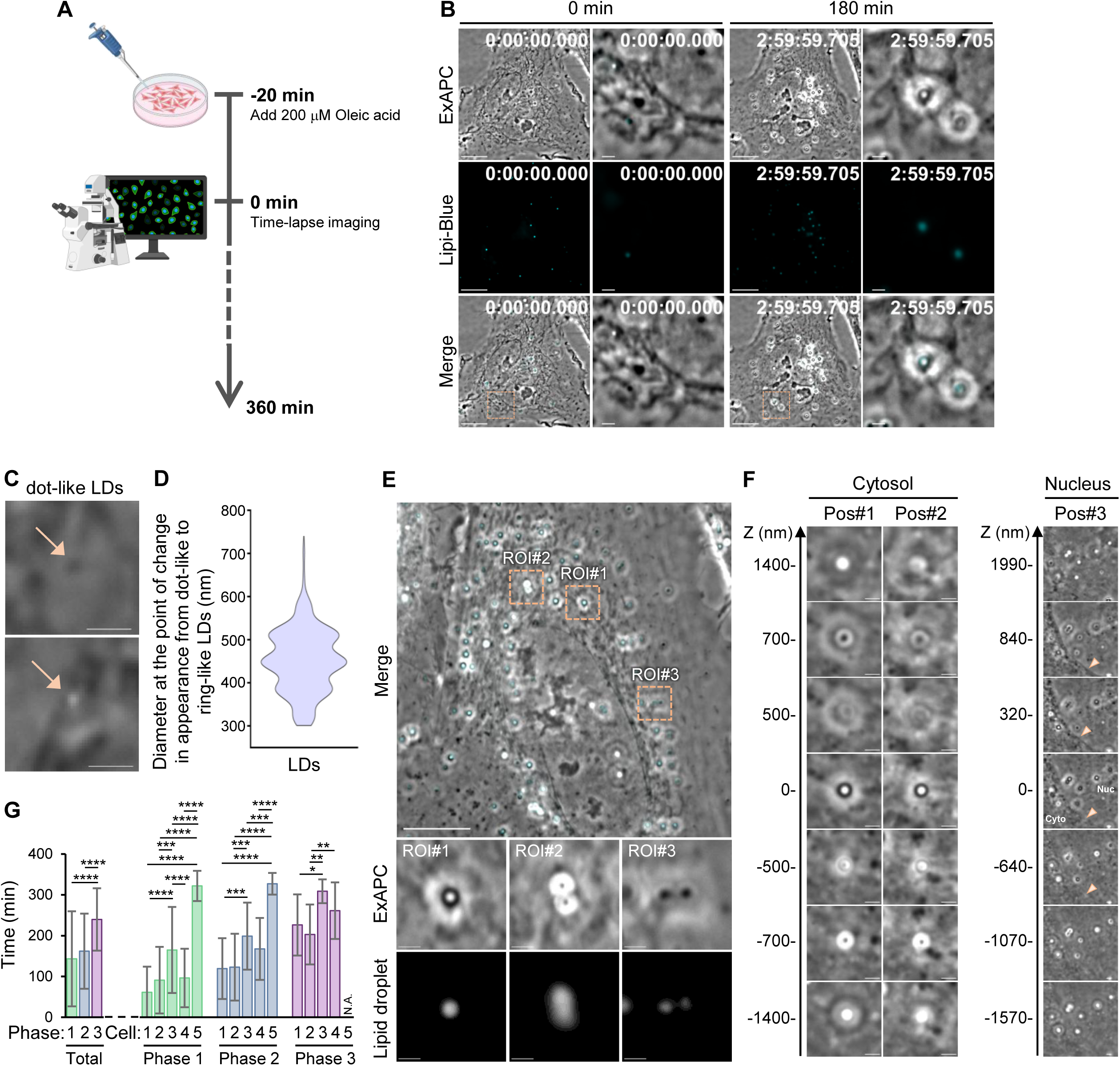
Visibility of LDs using ExAPC microscopy. (A) Schematic diagram of this experiment. (B) Oleic acid treatment-induced LD formation. HeLa cells were treated with oleic acid as described in Figure S3A. The cells shown here are the same as in Figure 5a. Scale bar: 10 μm; scale bar in inset: 1 μm. (C) Representative images of LDs in Phase 1. Scale bar: 1 μm. (D) Diameter at the point of change in appearance from dot-like to ring-like LDs (n = 165). (E) Representative images of LDs in Phases 2 and 3 in oleic acid-treated HeLa cells. (F) The visibility of LDs according to the difference in focal planes. Scale bar: 1 μm. (G) Time for each LD to reach each phase defined by diameter. Mean ± standard deviation. [Total] Phase 1: n = 176, Phase 2: n = 162, Phase 3: 123, [Phase 1] Cell#1: n = 27, Cell#2: n = 36, Cell#3: n = 42, Cell#4: n = 42, Cell#5: n = 29, [Phase 2] Cell#1: n = 33, Cell#2: n = 42, Cell#3: n = 36, Cell#4: n = 42, Cell#5: n = 9, [Phase 3] Cell#1: n = 25, Cell#2: n = 37, Cell#3: n = 8, Cell#4: n = 53, Cell#5: n = 0. One-way ANOVA for multiple comparisons was performed between phases (total) and between cells (phases 1-3).

**Figure S4.**
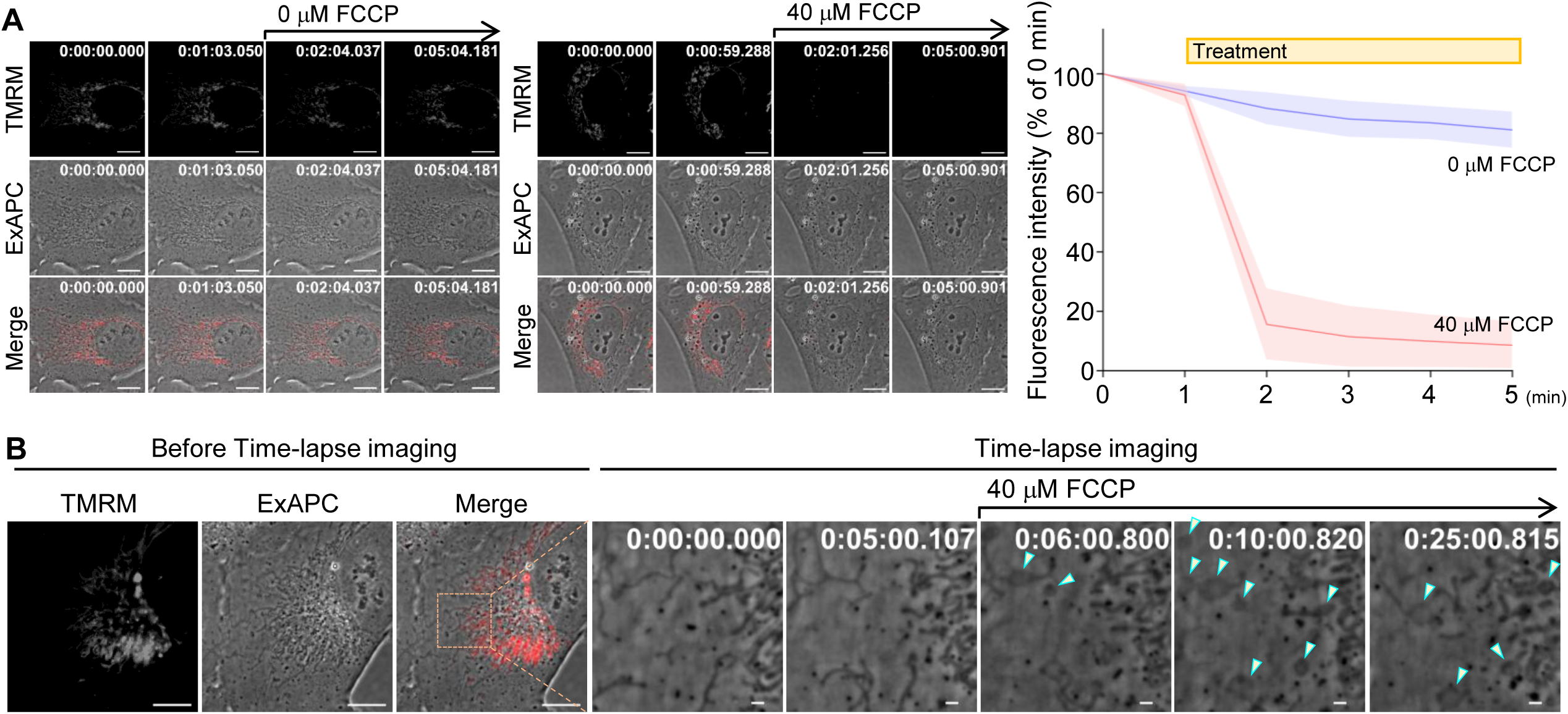
Mitochondrial dynamics in the presence of FCCP in U-2 OS cells. (A) Mitochondrial membrane potential in U-2 OS cells. Representative images at each time point are shown. All the data are presented as the mean ± standard deviation. 0 μM FCCP: n = 9 cells. 40 μM FCCP: n = 30 cells. (B) FCCP treatment-induced changes in mitochondrial morphology in U-2 OS cells. FCCP was added after image acquisition at frame 301 (1 frame/sec). Green arrowhead: Mitochondrion with morphological changes. Scale bar: (Before Time-lapse imaging): 10 μm, (Time-lapse imaging): 1 μm.

### Video 1. Sub-second time-lapse imaging

U-2 OS cells were imaged by time-lapse ExAPC microscopy to visualize the imaging intervals for a sub-second. Images acquired every 50 milliseconds, Video presented at 20 frames per second. Scale bar: 2.5 μm.

### Video 2. Cell division (HeLa cells)

HeLa cells were imaged by time-lapse ExAPC microscopy to visualize cell division. Images acquired every 1 minute, Video presented at 10 frames per second. Scale bar: 10 μm.

### Video 3. Cell division (iPSCs)

iPSCs were imaged by time-lapse ExAPC microscopy to visualize cell division. Images acquired every 1 minute, Video presented at 10 frames per second. Scale bar: 10 μm.

### Video 4. Apoptosis

HeLa cells were imaged by time-lapse ExAPC microscopy to visualize apoptosis. HeLa cells were treated with 1 μM STS at frame 15. Images acquired every 5 minutes, Video presented at 10 frames per second. Scale bar: 10 μm.

### Video 5. Entosis

ZR75-1 cells were imaged by time-lapse ExAPC microscopy to visualize entosis. Images acquired every 1 minute, Video presented at 10 frames per second. Scale bar: 10 μm.

### Video 6. BCLs

HeLa cells were imaged by time-lapse ExAPC microscopy to visualize BCLs. Images acquired every 1 second, Video presented at 10 frames per second. Scale bar: 10 μm.

### Video 7. IRS-1 droplet

U-2 OS cells expressing GFP-IRS1 were imaged by time-lapse ExAPC microscopy to visualize IRS-1 droplet. Images acquired every 1 minute, Video presented at 10 frames per second. Scale bar: 10 μm.

### Video 8. LD dynamics

Oleic acid-treated HeLa cells were imaged by time-lapse ExAPC microscopy to visualize LD dynamics. Images acquired every 20 seconds, Video presented at 10 frames per second. Scale bar: 1 μm.

### Video 9. Mitochondria fission

Hep3B cells were imaged by time-lapse ExAPC microscopy to visualize mitochondria fission. Images acquired every 1 second, Video presented at 10 frames per second. Scale bar: 1 μm.

### Video 10. Mitochondria fusion (end-to-end)

Hep3B cells were imaged by time-lapse ExAPC microscopy to visualize end-to-end fusion of mitochondria. Images acquired every 1 second, Video presented at 10 frames per second. Scale bar: 1 μm.

### Video 11. Mitochondria fusion (end-to-side)

Hep3B cells were imaged by time-lapse ExAPC microscopy to visualize end-to-side fusion of mitochondria. Images acquired every 1 second, Video presented at 10 frames per second. Scale bar: 1 μm.

### Video 12. FCCP treatment-induced changes in mitochondrial morphology in Hep3B cells

Hep3B cells were imaged by time-lapse ExAPC microscopy to visualize morphological changes of mitochondria in the presence of FCCP. Hep3B cells were treated with 40 μM FCCP at frame 301. Images acquired every 1 second, Video presented at 10 frames per second. Scale bar: 2.5 μm.

### Video 13. FCCP treatment-induced changes in mitochondrial morphology in U-2 OS cells

U-2 OS cells were imaged by time-lapse ExAPC microscopy to visualize morphological changes of mitochondria in the presence of FCCP. U-2 OS cells were treated with 40 μM FCCP at frame 301. Images acquired every 1 second, Video presented at 10 frames per second. Scale bar: 2.5 μm.

